# A disinhibitory circuit mechanism explains a general principle of peak performance during mid-level arousal

**DOI:** 10.1101/2023.07.28.550956

**Authors:** Lola Beerendonk, Jorge F. Mejías, Stijn A. Nuiten, Jan Willem de Gee, Johannes J. Fahrenfort, Simon van Gaal

## Abstract

Perceptual decision-making is highly dependent on the momentary arousal state of the brain, which fluctuates over time on a scale of hours, minutes, and even seconds. The textbook relationship between momentary arousal and task performance is captured by an inverted U-shape, as put forward in the Yerkes-Dodson law (Yerkes and Dodson, 1908). This law suggests optimal performance at moderate levels of arousal, and impaired performance at low or high arousal levels. However, despite its popularity, the evidence for this relationship in humans is mixed at best. Here, we use pupil-indexed arousal and performance data from various perceptual decision-making tasks to provide converging evidence for the inverted U-shaped relationship between spontaneous arousal fluctuations and performance across different decision types (discrimination, detection) and sensory modalities (visual, auditory). To further understand this relationship, we built a neurobiologically plausible mechanistic model and show that it is possible to reproduce our findings by incorporating two types of interneurons that are both modulated by an arousal signal. The model architecture produces two dynamical regimes under the influence of arousal: one regime in which performance increases with arousal, and another regime in which performance decreases with arousal, together forming an inverted U-shaped arousal-performance relationship. We conclude that the inverted U-shaped arousal-performance relationship is a general and robust property of sensory processing. It might be brought about by the influence of arousal on two types of interneurons that together act as a disinhibitory pathway for the neural populations that encode the available sensory evidence used for the decision.

**Significance statement:** When people are repeatedly performing the exact same task, their performance fluctuates over time. One of the causes of this behavioral variability is spontaneous fluctuations in arousal. According to the Yerkes-Dodson law, task performance is optimal at moderate levels of species’ arousal, with impaired performance at very low or high arousal levels. However, until now, the evidence supporting this law as a general mechanism in human decision-making is mixed, and a neural mechanism that may explain the inverted U-shaped arousal-performance relationship is lacking. We show that the Yerkes-Dodson law is a general law that holds for human observers across decision-making tasks and settings. Furthermore, we present a simple and neurobiologically plausible mechanistic model that can explain its existence.

## Introduction

Repeated presentations of the same noisy sensory input often lead to varying perceptual decisions (Rahnev & Denison, 2018; Wyart & Koechlin, 2016). One of the factors that contributes to this behavioral variability is the arousal state of the brain, which fluctuates spontaneously over time, causing variability in neuronal responsiveness to sensory stimulation (also often referred to as brain state) (Greene et al., 2023; McCormick et al., 2020; Waschke et al., 2021). The main drivers of these arousal fluctuations are the catecholaminergic and cholinergic neurotransmitter systems originating from the locus coeruleus and the basal forebrain, respectively. These neuromodulators have widespread effects throughout the brain, from controlling global cortical dynamics (e.g., low frequency power (McGinley, Vinck, et al., 2015)) to tuning the gain of sensory processing (McGinley, David, et al., 2015; Reimer et al., 2014; Vinck et al., 2015). Pupil dilation reflects the activity of neuromodulatory nuclei and can therefore be used to non-invasively approximate fluctuations in arousal state (de Gee et al., 2017; Joshi et al., 2016; Lloyd et al., 2023; Murphy, O’Connell, et al., 2014; Reimer et al., 2016; Strauch et al., 2022).

Although changes in arousal state affect decision-making abilities in both humans and mice (Murphy, Vandekerckhove, et al., 2014; Neske et al., 2019; Podvalny et al., 2021; van Kempen et al., 2019; Waschke et al., 2019; Xu et al., 2023), it remains unclear which level of arousal supports optimal performance. In mice, decision-making seems optimal at intermediate levels of arousal (McGinley, David, et al., 2015; McGinley, Vinck, et al., 2015), reflecting an inverted U-shaped relationship, in line with the well-known Yerkes-Dodson law (Yerkes & Dodson, 1908). In humans however, evidence is mixed. While some studies indeed show U-shaped arousal-performance associations in humans (Bullock et al., 2017; Murphy et al., 2011; Neske et al., 2019; Podvalny et al., 2021; van Kempen et al., 2019; Waschke et al., 2019), others report linear associations (Gelbard-Sagiv et al., 2018; Murphy, Vandekerckhove, et al., 2014; Podvalny et al., 2021).

Here we addressed these mixed results in the (human) literature and aimed to improve our understanding of the neural mechanisms underlying the arousal-performance association, which is not well understood in any species. To do so, we collected a large data set containing six behavioral tasks that differed in terms of the decision type (detection versus discrimination response) and the sensory modality being recruited (audition versus vision) using a within-subject design (**Figure 1A**). This dataset allowed us to explore the shape of the relationship between naturally occurring spontaneous fluctuations in pre-stimulus pupil-linked arousal and perceptual decision-making in humans. Aside from different sensory modalities and decision types, all tasks had comparable trial structures and behavioral accuracy was matched across all tasks (see Methods for details). To anticipate our findings, we observed that the arousal-performance relationship was clearly inverted U-shaped, regardless of the sensory modality or decision type at hand.

**Figure 1.**
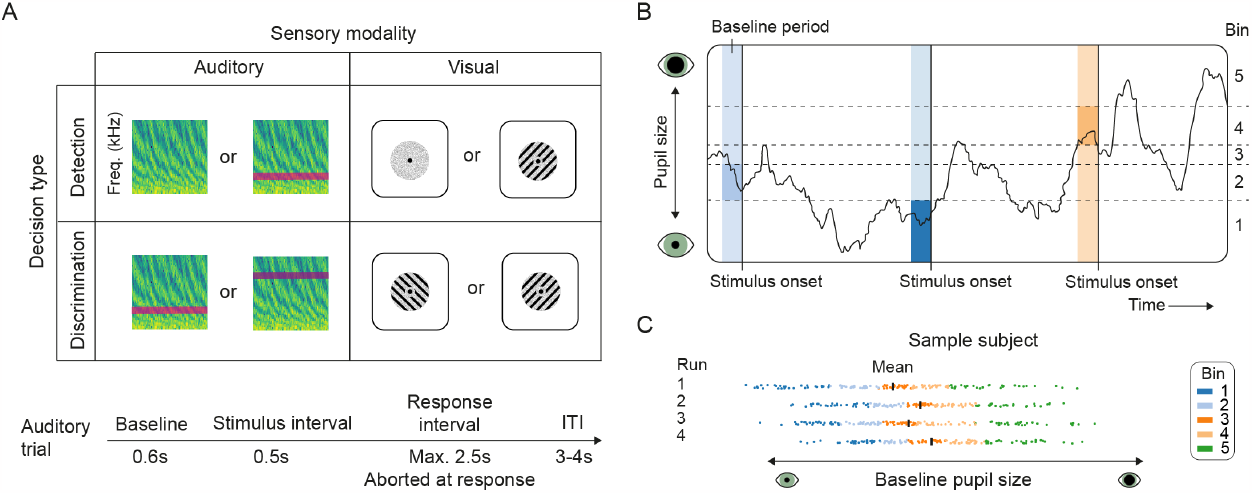
Behavioral tasks and binning analysis. (A) General task overview. (B) Example of the continuous pupil signal of three trials with baseline windows [-.5-0s] divided over five bins. Shaded areas indicate the baseline interval that was used to calculate baseline pupil size. (C) Example data of one session of the auditory detection task of a sample participant that shows that bins were determined for each task run (block) separately. Black vertical lines indicate the mean baseline pupil size of each depicted run. Each dot represents the baseline pupil size of a trial, and the color of the dots indicates bin membership (five bins).

After having established a robust inverted U-shaped relationship between spontaneous pupil-linked arousal fluctuations and task performance, and given that inverted U-shaped relationships have been observed in both mice and humans (Bullock et al., 2017; McGinley, David, et al., 2015; McGinley, Vinck, et al., 2015; Murphy et al., 2011; Neske et al., 2019; Podvalny et al., 2021; van Kempen et al., 2019; Waschke et al., 2019), we aimed to increase our understanding of the arousal-performance relationship by developing a minimal network model reproducing the observed results. We reasoned that explanations that rely on biological constraints common to both species are most prominent candidates to explain its underlying mechanisms. Optimal signal detection properties have been linked with maintaining a certain excitation-inhibition balance in neural circuits (Joglekar et al., 2018; Rubin et al., 2017; Tian et al., 2020; Vogels & Abbott, 2009) and in mice, a balanced competition between inhibition and excitation (or rather, disinhibition) has been found in circuits with pyramidal neurons and inhibitory cells expressing parvalbumin (PV), somatostatin (SST) and vasoactive intestinal peptide (VIP) (Garcia Del Molino et al., 2017; Litwin-Kumar et al., 2016; Pakan et al., 2016; C. K. Pfeffer et al., 2013; Rudy et al., 2011). As such circuitry is also present in human and non-human primates (Kooijmans, Sierhuis, et al., 2020; Kooijmans, Upschulte, et al., 2020), mechanisms based on this inhibition/disinhibition balance could provide an explanation for the observed phenomenon across species.

To provide a mechanism that might explain the inverted U-shaped arousal-performance relationship in our detection and discrimination tasks, we built a minimal computational model, adapted from well-known population-based models of neural dynamics used to describe performance on decision-making tasks (Wong & Wang, 2006). Our model describes in detail the temporal evolution of the global synaptic conductances corresponding to NMDA and GABA receptors of two competing excitatory populations that are indirectly modulated by an arousal signal. Based on our computational model, we conclude that the inverted U-shaped arousal-performance relationship might result from the influence of arousal on two types of interneurons that together act as a disinhibitory pathway for the neural populations that encode sensory evidence.

## Results

Participants (N=28) took part in three all-day (9am-4pm) experimental sessions in which six different tasks were performed (see Methods for details). In brief, in the auditory domain, participants either discriminated between high and low pitch tones in one task or they detected a low pitch tone vs noise in another (present/absent judgement). In the visual domain, two versions of a visual Gabor detection task and two versions of a visual Gabor orientation discrimination task were performed. Trials of all tasks followed a similar structure with a baseline period, stimulus interval, response interval, and a variable inter trial interval (**Figure 1A**), although visual tasks were generally faster paced than auditory tasks (see Methods for details). All tasks were titrated to reach 75% accuracy (the mean accuracy across tasks was 76.3%). First, we tested whether the overall relationship between pupil-indexed arousal and perceptual decision-making was linearly or U-shaped, irrespective of specific task parameters. To this end, we combined the data of all six behavioral tasks, harnessing the large statistical power of our complete dataset (3570 trials per participant on average). This analysis across tasks gives us a general indication of the arousal-performance relationship, irrespective of sensory modality and decision type. For each run separately, the trials were distributed across twenty equally populated bins based on the average pupil size in the pre-stimulus baseline period (500ms window preceding the target, **Figure 1B-C**). Note that all bins contained equal numbers of trials, and that therefore the bins did not have equal widths (**Figure 1B-C**). Next, we calculated Signal Detection Theoretic sensitivity (SDTs d’ (Green & Swets, 1966)) and average reaction time (RT) for each pupil bin across all tasks. We used linear mixed models and formal model comparison to assess whether the arousal-performance relationship was linear or quadratic. Using both AIC (Akaike information criterion (Akaike, 1974)) and BIC (Bayesian information criterion (Schwarz, 1978)) we directly compared evidence for linear versus quadratic models. A difference in AIC or BIC values of more than 10 is considered evidence for the winning model to capture the data significantly better (Burnham & Anderson, 2004). In our case, both ΔAIC and ΔBIC were strongly in favor of a quadratic relationship between pre-stimulus pupil size and sensitivity (ΔAIC=31.0, ΔBIC=26.7) and RT (ΔAIC=20.3, ΔBIC=16.0). In other words, the overall arousal-performance relationship was inverted U-shaped, with highest sensitivity and shortest RTs at intermediate levels of pupil-linked arousal (**Figure 2**, ‘All’). The association between pupil size and sensitivity was much more pronounced than the association between pupil size and RT, which becomes especially clear when both performance measures are expressed in percent signal change (**Supplementary Figure 1**).

**Figure 2.**
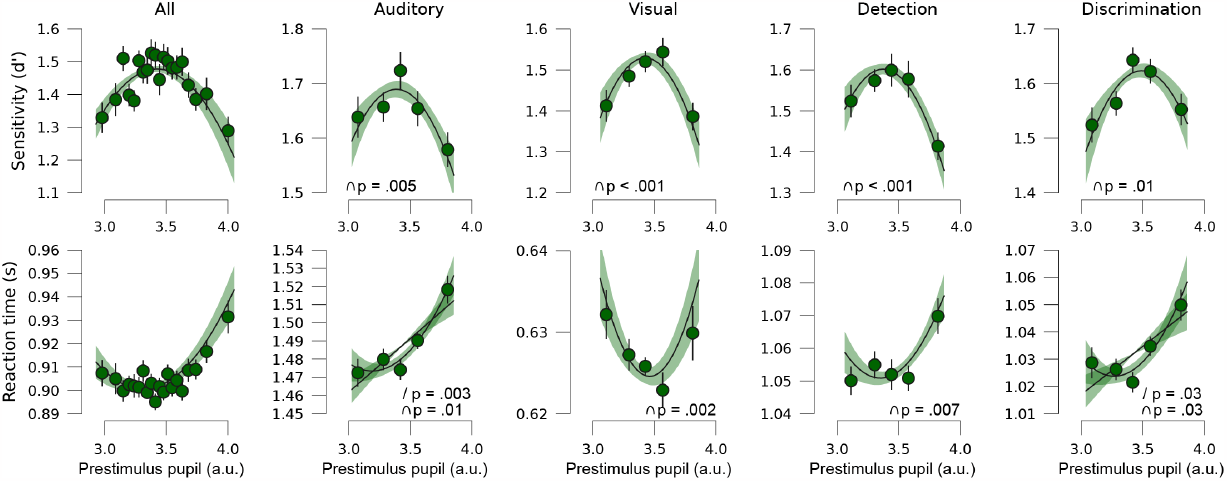
Perceptual decision-making is optimal at moderate levels of pupil-linked arousal across sensory modalities and decision types. Perceptual sensitivity (d’, top row) and RT (bottom row) for small to large baseline pupils for all tasks combined, auditory tasks, visual tasks, detection tasks, and discrimination tasks, from left to right respectively. We show polynomial regression lines for first and second order fits when significantly different from zero (average regression line of all participants, shading reflects standard error of the mean (SEM), see Methods). Error bars on the data points represent SEM of the within-subject variation across tasks.

We performed several control analyses to further understand these results. First, it has been argued that the (quadratic) relationship between pupil-linked arousal and performance may be driven by time on task (van den Brink et al., 2016), but we found no evidence for this. After regressing out the influence of time on task on pre-stimulus pupil size, the quadratic relationship remained intact (see Methods and **Supplementary Figure 2**). Next, we performed three additional analyses to test whether the quadratic arousal-performance relationship was driven by significant behavioral or pupil events on the previous trial. Specifically, 1) we excluded all trials that followed erroneous responses (23.1% of all trials), possibly upregulating arousal, 2) we performed the analyses after regressing out task-evoked pupil responses on the previous trial (de Gee et al., 2014, 2022; Waschke et al., 2021), and 3) we tested whether the quadratic arousal-performance relationship would also be revealed if the task-evoked pupil response in the previous trial was used as the measure of the pupil-linked arousal state. These analyses further solidified our main conclusions and indicated that the inverted U-shaped relationship between pre-stimulus (baseline) pupil size was not driven by response errors or task-evoked pupil responses on the previous trial (**Supplementary Figure 3**). Taken together, these control analyses indicate that the inverted U-shaped arousal-performance relationship described here (**Figure 2**) is likely driven by spontaneous fluctuations in pupil-indexed arousal.

Next, we addressed whether the arousal-performance relationship depends on the decision type (detection vs discrimination) or the sensory modality (visual vs auditory). To test this, we performed polynomial regression for each sensory modality and decision type separately, averaging across tasks that fell into each category (see Methods for details). This allowed us to step aside from specific experimental manipulations of each of the six tasks individually, which will be reported elsewhere (in the future) (see e.g., (Nuiten et al., 2023), and focus on generalities across these tasks, namely which sensory modality was recruited (visual or auditory) and what was the task to be performed (discrimination or detection). Additionally, this ensured enough trials per analysis to reliably compute regressions across pupil bins. To further warrant statistical power, these analyses of subsets of the data were performed with five pupil size bins to account for lower trial numbers after splitting the data (**Figure 1B and C**).

Similar to the first analysis, perceptual sensitivity (d’) was highest and RT lowest at intermediate levels of pupil-linked arousal for both auditory and visual tasks (auditory d’ ß_2_: t(26)=-2.79, p=.005, d=-0.55, visual d’ ß_2_: t(27)=-3.88, p<.001, d=-0.75; auditory RT ß_2_: t(26)=2.41, p=.01, d=0.47; visual RT ß_2_: t(27)=3.26, p=.002, d=0.63), and for both detection and discrimination tasks (detection d’ ß_2_: t(27)=-3.96, p<.001, d=-0.76, discrimination d’ ß_2_: t(27)=-2.39, p=.01, d=-0.46; detection RT ß_2_: t(27)=2.65, p=.006, d=0.51, discrimination RT ß_2_: t(27)=2.05, p=.03, d=0.39; **Figure 2**). Again, the association between pupil size and sensitivity was much more pronounced than the association with reaction times (see **Supplementary Figure 1**). Note that two out of four RT analyses additionally showed positively linear relationships (auditory RT ß_1_: t(26)=3.29, p=.003, d=0.70; discrimination RT ß_1_: t(27)=2.30, p=.03, d=0.43), further elaborated on in the Discussion. Interestingly, the pupil-sensitivity relationship appeared to be mainly driven by an effect of pre-stimulus pupil size on hit rate, rather than false alarm rate (**Supplementary Figure 4**), in line with observations in mice (McGinley, David, et al., 2015; McGinley, Vinck, et al., 2015). In sum, the inverted U-shaped arousal-performance relationship can be observed for both decision types and for both sensory modalities (most prominently for d’), despite general differences in the overall timing of these tasks (visual tasks were faster paced). Therefore, optimal performance at moderate levels of spontaneous pupil-linked arousal appears to be a general and robust characteristic of perceptual decision-making.

Even though the inverted U-shaped arousal-performance relationship has been reported before, it remains unclear how it can emerge from neural interactions. In principle, inverted U-shapes would require a change in the dynamical regime of the underlying cortical circuits, and these are not well known. One plausible candidate mechanism is the interplay between different interneuron types expressing parvalbumin (PV), somatostatin (SST) and vasoactive intestinal peptide (VIP), since it seems to play a role in disinhibitory and paradoxical dynamics (defined as a counterintuitive response of a circuit to a given external input) in cortical circuits (Dipoppa et al., 2018; Garcia Del Molino et al., 2017; Lee et al., 2013; Pakan et al., 2017; Tsodyks et al., 1997). We therefore explore whether the interplay between PV, SST and VIP can explain the nonlinear responses to arousal input observed.

We built a firing rate-based computational model of a cortical circuit performing a detection and/or discrimination task (Wong & Wang, 2006) under the influence of a hypothetical arousal signal that is highly correlated with pupil size. The circuit describes the continuous temporal evolution of biophysical variables –such as firing rates and synaptic conductances of neural populations (see Methods for more details) and describes the interactions between two excitatory neural populations (E_A_ and E_B_; encoding the two behavioral choices A and B) mediated by synapses with NMDA receptors (**Figure 3A**). The model incorporates a non-selective population of PV interneurons, whose role is to provide a baseline level of inhibition and to mediate the competition between both excitatory populations. In the absence of any modulatory signal or further components, this constitutes a well-studied model (Wong & Wang, 2006) which reflects competitive dynamics between choices A and B. More precisely, both excitatory populations receive sensory input, which propels them to ramp up their own activity and suppress the activity of the other excitatory population –a process mediated by the inhibitory population PV. This gives rise to a winner-take-all decision process (Wong & Wang, 2006), which corresponds in our case to a detection (A: present, B: absent) or a discrimination task (choice A, choice B). A decision is made when the firing rate of one of the excitatory populations reaches a certain predefined threshold (**Figure 3B**).

**Figure 3.**
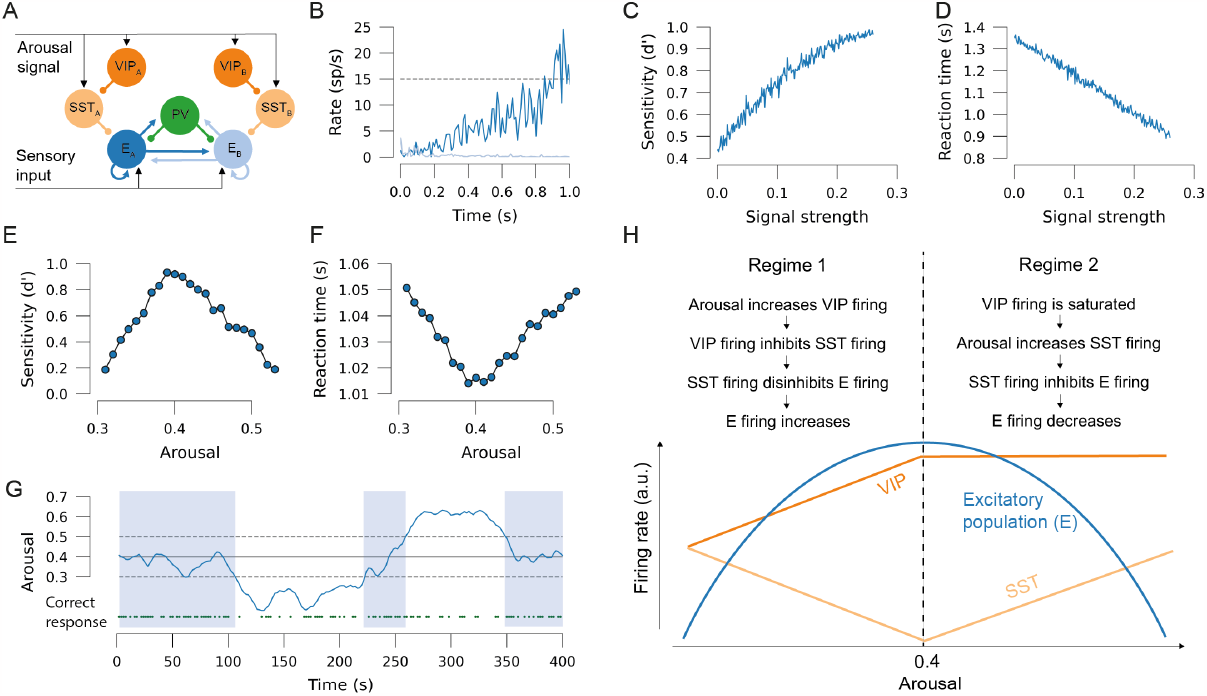
Computational model explaining decision-making behavior. (A) Schematic description of the model, with excitatory populations E_A_ and E_B_ encoding two possible choices in a decision-making task (choice A and B), which is also modulated by a common arousal signal to VIP and SST cells, which are also specific to choice A or B. Connectivity follows electrophysiological evidence and previous modeling approaches. Arrows indicate excitatory connections, dots indicate inhibitory connections. (B) Example of the excitatory activity of a decision, in which population E_A_ (dark blue) wins over population E_B_ (light blue) when its firing rate reaches the decision threshold. (C) Performance of the model, defined as the sensitivity (d’) for 300 trials, as a function of the normalized input, signal strength. (D) Mean RT of the model as a function of input, signal strength. (E) Performance of the model as a function of the strength of the arousal signal. (F) Mean RT versus arousal strength. Results in E-F averaged over 3000 trials. (G) Simulation of a ∼400 second session, corresponding to 266 consequtive trials, in which the level of arousal (blue line) slowly fluctuates over time and modulates the probability of making a correct choice (green dots). The optimal arousal level is indicated with a solid horizontal line. Dashed lines indicate standard deviation of the arousal signal and shaded areas indicate that arousal is within one standard deviation of the optimal arousal level. (H) Schematic description of how the interaction between arousal, VIP and SST cells brings about the inverted U-shaped relationship between arousal and perceptual decision-making. For panel B-D, the arousal level is 0.4. For panels B and E-G, the signal strength is 0.1.

To incorporate the effects of arousal modulation, which were not considered in previous work (Wong & Wang, 2006), our model here also incorporates populations of SST and VIP cells, one of each per behavioral choice, as these cells have been found to be choice-specific (D. Kim et al., 2016). Following experimental evidence, VIP and SST cells form a connectivity motif which may disinhibit the activity of the corresponding excitatory populations (C. K. Pfeffer et al., 2013) and receive top-down input from other brain areas. We assumed that these neural populations act as a disinhibitory pathway to excitatory cells (C. K. Pfeffer et al., 2013), although we also considered that the dynamics of the VIP-SST-excitatory cell circuit can switch from disinhibition to inhibition depending on the input level and other conditions (Garcia Del Molino et al., 2017). Finally, we considered that sensory input was delivered to the excitatory populations and that a hypothetical arousal signal reached both VIP and SST populations (similar to a top-down modulatory signal).

An example trial of the model in the absence of any modulation by the arousal signal, and therefore equivalent to the situation modelled in previous work (Wong & Wang, 2006), is shown in **Figure 3B**, where the competition between both excitatory populations is won by choice A as the corresponding firing rate E_A_ reaches the threshold of 15 spikes/s. To verify that the model showed basic detection behavior, we first varied the signal strength of the sensory input. Sensitivity and chronometric curves of the model output choices (**Figure 3C and D**), reflected that input detections became easier and were quicker for trials with higher signal strength, similar as the case for decision-making tasks (Roitman & Shadlen, 2002) and in agreement with previous models (Wong & Wang, 2006).

When we simulated the model for different strengths of the arousal signal arriving to SST and VIP populations, while keeping signal strength constant as in the human experiments, we observed that the sensitivity of the model followed an inverted U-shape with respect to the arousal level (**Figure 3E**). In the model, this behavior emerges due to the existence of two dynamical regimes, one for low arousal levels (i.e., arousal strength smaller than 0.4), and another for high arousal levels (larger than 0.4) (**Figure 3H**; also see **Supplementary Figure 6**). For low arousal levels, the system is typically in a suboptimal state –in the sense that the firing rates evoked by the sensory input are rarely elevated enough to trigger a detection decision. As the arousal level increases, the VIP activity inhibits the firing of SST cells, which in turn disinhibits the excitatory populations. This provides an additional input, which makes it easier for the circuit to correctly detect the sensory signal, and the sensitivity overall increases. This trend is maintained until arousal levels of ∼0.4 are reached in the model, corresponding to optimal detection, when VIP cells are strongly driven by the arousal signal and high or even saturated firing rate levels of VIP firing are reached. For higher arousal levels, VIP inhibition to SST is not able to compensate for the excitatory effects of the arousal signal to SST. Therefore, the net effect of the arousal signal in SST cells now becomes excitatory, leading to the inhibition of the excitatory populations. This leads to a decrease in performance with further increasing arousal levels as the circuit returns to a suboptimal state, thus completing the inverted U-shape. See **Supplementary Figure 6** for a graphical depiction of the inhibitory strengths provided by SST and VIP at each of these conditions.

As easier trials (i.e., with stronger sensory input) lead to quicker reaction times in the model, we also obtained the corresponding U-shaped relationship between reaction times and arousal levels (**Figure 3F**) that we also found experimentally. In a realistic setting in which the arousal level would slowly fluctuate across trials within a given session, this mechanism would lead to changes in the detection probability across the session (**Figure 3G**). Due to the symmetry that the model assumes between excitatory populations A and B (**Figure 3A**), the results of the model are the same for the case of a discrimination task, in which the model must react to input for either excitatory population A or B.

## Discussion

We showed that perceptual variability of repeated identical near-threshold stimuli was predicted by ongoing fluctuations in cortical arousal, as indexed by pupillometry. More specifically, we demonstrated that the arousal-performance relationship is inverted U-shaped, with optimal performance at moderate levels of pupil-indexed arousal. This nonlinear relationship does not appear to depend on the sensory modality (vision/audition) or decision type (discrimination/detection) at hand and were also robust to potential differences in task-related factors, such as time on task (i.e., fatigue), potential upregulation of arousal by decision errors, task-evoked pupil responses on the previous trial, and general differences in the overall timing of these tasks. Therefore, the inverted U-shaped arousal-performance relationship appears to reflect a general, flexible, and robust property of sensory processing in humans.

To provide a potential neural mechanism explaining these observations, we built a simple computational model that incorporated different types of interneurons, VIP and SST, which are both modulated by an arousal signal. In the case of humans and other primates, similar action may be expected from interneurons expressing calbindin (CB) and calretinin (CR), which are thought to be analogous to SST and VIP in rodents (Medalla et al., 2023) and similarities between both circuits have been well documented (Kooijmans, Sierhuis, et al., 2020). When we simulated different arousal strengths, this architecture produced two dynamical regimes that depend strongly on the firing rate of the interneurons. In the first regime, VIP firing increases under the influence of increasing arousal, thereby inhibiting SST firing and disinhibiting the firing of the excitatory populations, leading to increasing performance with increasing arousal. Conversely, in the second regime, performance decreases with increasing arousal as a result of saturated VIP firing, disinhibition of SST firing, and subsequent inhibition of the excitatory populations (**Figure 3H**). Together, these regimes form the inverted U-shaped relationship between arousal and perceptual decision-making that we observed in our empirical data.

The computational model proposed to explain the observed experimental findings constitutes only a first step towards a complete understanding of the mechanisms underlying the Yerkes-Dodson law. The model also relies on several hypotheses, which will need to be tested in future experimental studies, although some hypotheses are already partially supported by existing data. For example, our model requires the presence of a disinhibitory mechanism involving SST and VIP cells. It is unclear which brain region is responsible for the inverted U-shape relationship, but SST and VIP cells are found throughout the neocortex (Y. Kim et al., 2017) and such disinhibitory mechanisms have been found across multiple areas, including visual and somatosensory cortex (Dipoppa et al., 2018; Lee et al., 2013; C. K. Pfeffer et al., 2013). We also assumed that arousal signals should be able to modulate the firing of SST and VIP cells, which is plausible given that these cell types are known to be modulated by top-down signals (C. K. Pfeffer et al., 2013). Finally, the saturation of VIP activity is crucial for our mechanism to replicate the inverted U-shape. Electrophysiological and optogenetics studies have shown that VIP cells indeed display spike frequency adaptation (Badrinarayanan et al., 2021; Canto-Bustos et al., 2022), which should be able to limit the response of VIP cells receiving large arousal signals. In addition, strong short-term synaptic depression effects have been found in GABAergic cells in layer 2/3 (Lombardi et al., 2023), where VIP cells are the most common interneuron type (Billeh et al., 2020; Rudy et al., 2011). Previous computational work has shown that the combination of spike frequency adaptation (or other types of neuron-level adaptation) and short-term synaptic depression is highly effective to drastically reduce signal transmission and generate saturation of activity (Mejías & Torres, 2008; Mejias & Torres, 2011; Torres et al., 2011), suggesting that VIP saturation is a biologically plausible scenario in this context.

More precise descriptions of the influence of arousal signals on task performance will be necessary to extend and improve our current model in the future. For example, it is well known that acetylcholine and catecholamines have a wide range of modulatory effects on task performance, which are not explicitly included in our present model. Likewise, we have focused here on the effects of arousal signals on SST and VIP cells, which constitutes a simplification. In this approach we do not consider how arousal-related signals, such as cholinergic modulation for example, affect the physiological properties of pyramidal cells, such as excitability (Cole & Nicoll, 1984a, 1984b), spike frequency adaptation (Barkai & Hasselmo, 1994; Madison & Nicoll, 1986a, 1986b) and modulation of glutamatergic transmission (Hasselmo, 2006). Future modeling work should aim to incorporate some of these properties and investigate their potential effects on the Yerkes-Dodson law.

Besides providing a mechanistic explanation for our experimental results, our computational model also makes some testable predictions. As these predictions involve the activity of VIP and SST cells, we propose that optogenetic techniques used in mice performing tasks during different arousal conditions (Klaver et al., 2023) would be a potential way to validate these predictions experimentally. First, we predict that inhibitory and disinhibitory dynamics mediated by SST and VIP cells play a fundamental role in the emergence of optimal levels of arousal for decision-making tasks, and interfering with these mechanisms should abolish (or modify) the inverted U-shaped relationship. Second, we anticipate that VIP firing will be moderate for low arousal levels and, due to VIP saturation, will be elevated and similar for both intermediate and high arousal levels. Likewise, SST will display moderate firing rates for low and high arousal levels, and low firing rates for intermediate arousal levels. We expect that further development of this model will provide additional insight into the role of arousal in detection and discrimination tasks, for example by extending this model to include more cortical areas – in line with recent advances (Klaver et al., 2023; Mejías & Wang, 2022).

Our results bear a strong resemblance to various findings of an inverted U-shaped relationship between pre-stimulus oscillatory power dynamics in the 8-12 Hz (alpha) band and perceptual decision-making (Linkenkaer-Hansen et al., 2004; Zhang & Ding, 2010). Indeed, pupil size and pre-stimulus alpha power are believed to be two sides of the same coin: high arousal states go hand in hand with alpha power suppression and pupil dilation (Aston-Jones & Cohen, 2005; Reimer et al., 2016), and alpha power and pupil diameter appear to be coupled during quiet wakefulness (Montefusco-Siegmund et al., 2022). Yet, there also appear to be dissociations between the effects of pre-stimulus alpha power and pupil-linked arousal on behavior (Lloyd et al., 2023), and there is evidence that suggests that the relationship between alpha power and pupil size is nonlinear (T. Pfeffer et al., 2021; Podvalny et al., 2021). Moreover, a set of studies indicates that pre-stimulus alpha oscillations drive decision bias by modulating neuronal excitability (Iemi et al., 2017, 2022; Kloosterman et al., 2019; Samaha et al., 2020), rather than perceptual sensitivity as we report here. Future work should aim to bridge the gap between these parallel fields that investigate arousal through pre-stimulus pupil size and pre-stimulus oscillatory dynamics to further our understanding of the influence of cortical state on perceptual decision-making.

The relationship between arousal and sensitivity in our study was clearly inverted U-shaped across both sensory modalities and decision types. For reaction times, however, we observed additional linear relationships for two out of four task variations (note that arousal-RT associations were less strong than for d’ however). We cannot completely rule out the possibility that the arousal-RT relationship depends on sensory modality or decision type, and future work is needed to address the issue. In fact, although the observed quadratic arousal-sensitivity relationship was clearly robust to task parameters, some previous studies in humans have also reported linear effects. In mice, primarily U-shaped functions are observed (de Gee et al., 2022; McGinley, David, et al., 2015; McGinley, Vinck, et al., 2015): mice perform best at an auditory detection task during quiet wakefulness, and worse during states of low and high arousal (including locomotion) (McGinley, David, et al., 2015). In humans however, the shape of the arousal-performance relationship appears to be less clear. Some studies have suggested that increased arousal linearly improves perceptual decision-making (Gelbard-Sagiv et al., 2018; Murphy, Vandekerckhove, et al., 2014; Podvalny et al., 2021), whilst others report optimal performance at intermediate levels of arousal (Bullock et al., 2017; Murphy et al., 2011; Neske et al., 2019; Podvalny et al., 2021; van Kempen et al., 2019; Waschke et al., 2019). Although we have not been able to find clear consistencies in the available literature causing this dichotomy, we identify (at least) four possible explanations. First, it is possible that the studies that report linear effects employed easier tasks than we did here (or less perceptually complex tasks (Sörensen et al., 2022)). The seminal work by Yerkes and Dodson (1908) predicts a quadratic effect of arousal on performance for difficult tasks, but a (positive) linear effect for easy tasks. There is hardly any behavioral evidence for the difference in the shape of arousal-performance relationships between easy and difficult tasks in the domain of perception, but this issue deserves further investigation (Sörensen et al., 2022). Second, studies reporting linear effects might investigate different parts or only a limited range of the arousal spectrum. To illustrate, it may be that if one only samples the low end of the arousal spectrum (low to medium arousal), one may end up with a positive linear relationship between arousal and performance (hence missing the downward going part for high arousal states). Third, it is possible that the quadratic relationship that we report only holds for spontaneous fluctuations in arousal during task performance whilst sitting still – and not for other arousal levels, such as pharmacologically manipulated states (Gelbard-Sagiv et al., 2018) or arousal levels ranging from quiet wakefulness to locomotion (Bullock et al., 2017). Fourth, quadratic relationships are not always considered, for instance when only two arousal states are compared (Xu et al., 2023). This may relate to a potential lack of statistical power (e.g., number of trials) that is necessary to dissect the data in (enough) bins to properly uncover the quadratic relationship between arousal and decision behavior.

In conclusion, our findings attribute a general role to spontaneous arousal fluctuations in the modulation of perceptual decision-making. Independent of task parameters, behavioral performance appears to be optimal at intermediate levels of pupil-linked arousal. Based on a new computational model, we propose that this inverted U-shaped arousal-performance relationship might result from the influence of arousal on VIP and SST interneurons that together act as a disinhibitory pathway for the neural populations that encode the available sensory evidence.

## Methods

### Subjects

For this study, 30 right-handed Dutch speaking male participants (aged between 18-30) were recruited from the University of Amsterdam. Because this study involved a pharmacological manipulation, all participants underwent extensive medical and psychological screening to rule out any medical or mental illnesses. All participants gave written consent for participation and received monetary compensation for participation. This study was approved both by the local ethics committee of the University of Amsterdam and the Medical Ethical Committee of the Amsterdam Medical Centre. All participants provided written informed consent after explanation of the experimental protocol. Two participants decided to withdraw from the experiment after the first experimental session. The data from these participants has been excluded from further analyses, resulting in N=28. Due to technical malfunction, one participant did not perform the auditory tasks, resulting in N=27 for the auditory data.

### Procedure

Participants took part in three experimental sessions that were separated by at least one week each. On each session, participants received one of three drugs in a random and counterbalanced order: placebo, atomoxetine, or donepezil, but here we only analyze the placebo data. In the first two hours of each session, participants performed auditory discrimination (session 1 and 2) and detection tasks (session 3). In the second half of all sessions, participants performed four versions of visual detection and discrimination tasks (all sessions; the order of the tasks was randomized between participants).

After arriving at the testing site, participants received either placebo (2 sessions) or donepezil (1 session). Right after oral ingestion of the pharmaceutical, participants would perform an auditory staircase task. The staircase procedure lasted approximately 30 minutes and was immediately followed by the main auditory task, which lasted approximately 1.5 hours. After the auditory task, participants then received placebo (2 sessions) or atomoxetine (1 session) to ensure that both donepezil and atomoxetine plasma concentrations would peak at the start of the visual tasks (four hours after the start of the session). The pharmacological intervention is not of interest to this work; therefore, we only analyze the placebo data of the visual tasks. As donepezil plasma concentrations peak after approximately four hours, we assume that donepezil blood concentration levels were negligible during the auditory tasks (i.e., the first two hours after donepezil ingestion). For the auditory tasks, we thus analyze the data from all sessions. Below, we provide a brief description of the behavioral tasks, for more details on the pharmacological manipulation, visual tasks and additional results please see (Nuiten et al., 2023). During all tasks, participants were seated 80cm from a computer monitor (69x39cm, 60Hz, 1920x1080 pixels) in a darkened, sound isolated room. To minimize head movements, participants rested their heads on a head-mount with chinrest. All tasks were programmed in Python 2.7 using PsychoPy (Peirce, 2007) and in-house scripts.

### Behavioral tasks

#### Auditory detection and discrimination tasks

Participants performed two different auditory tasks on separate sessions. The first task was an auditory discrimination task (session 1 and 2), in which participants had to discriminate between a high pitch tone (50% of trials) and a low pitch tone (50% of trials) against a background of auditory noise (**Figure 1A**). The other task was an auditory detection task (session 3) in which participants had to indicate whether they deemed a target tone to be present or absent alongside auditory noise that was presented on every trial (50% target present trials; **Figure 1A**). Besides performing the discrimination or detection task, participants were asked to report the confidence (low or high) in each perceptual decision that they made (not analyzed here). The perceptual decisions and confidence reports were simultaneously given by pressing one of four buttons (*A, S, K, L*) on the keyboard. Because the auditory stimuli in these tasks do not inherently convey any direction (such as the visual stimuli discussed below), the meaning of the buttons with respect to the perceptual decisions was counterbalance across subjects. The buttons’ mapping to confidence was the same for all participants (*A* and *L* for confident, *S* and *K* for not confident).

The auditory stimulus set consisted of dynamic noise and two pure tones (sine waves). The dynamic noise (so-called temporally orthogonal ripple combinations; TORCs) (Atiani et al., 2009; McGinley, David, et al., 2015) was present on every trial and was always played at the same volume (61.1 dB). On top of the dynamic noise, there was a low pitch sine wave (300 Hz) and a high pitch sine wave (350 Hz) in the discrimination task and only the low pitch tone was used for the detection task. For the discrimination task, the mean tone volume was 3.22% of the TORC volume. For the detection task, the mean tone volume was 3.75% of the TORC volume. Auditory stimuli were binaurally presented through over-ear headphones. The volume of the tones was determined per subject by means of a staircase procedure aimed at 75% correct (see <u>Staircasing procedure</u>). To ensure that performance remained around 75%, the volume of the target tones was automatically adjusted between runs. If performance in a run deviated between 5-10% of target performance (75%), the volume of the tones was adjusted with 5% of the current tone volume on the following run. If performance deviated more than 10% from target performance, target volume was adjusted with 10%.

Each auditory trial started with a baseline interval (600ms), followed by the stimulus interval (500ms) in which the auditory noise was always presented (**Figure 1A**). Depending on the task at hand, a target tone was or was not presented for the entire duration of the stimulus interval. After the offset of the sounds, participants had 2.5 seconds to respond (response interval). The response interval was aborted at response or after 2.5 seconds if no response was given, follow by a variable inter trial interval (randomly drawn from a uniform distribution between 3 and 4 seconds).

Participants performed 560 trials of the auditory tasks per session, distributed over four runs of 140 trials that lasted approximately 12 minutes each. As participants performed the auditory discrimination task on session 1 and 2, and the auditory detection task only on session 3, we have collected double the number of trials for the auditory task (1120 trials). During the auditory tasks, participants viewed a computer screen with a grey background and a black fixation shape in the center (Thaler et al., 2013).

#### Visual tasks

Participants performed four visual tasks. Two of these were variations of a discrimination task, in which participants were asked to discriminate the orientation of a Gabor patch hidden in dynamic visual noise as being rotated clockwise (CW: 45°, 50% of trials) or counter-clockwise (CCW: -45°, 50% of trials; **Figure 1A**). The other two tasks were detection tasks in which participants had to indicate whether they believed a Gabor patch (CCW or CW) to be present or absent (50% present trials). These two detection tasks differed in terms of response bias manipulation (conservative versus liberal, see below). Besides these main task characteristics, there were some additional instructions and manipulations that are discussed below. During all visual tasks, gaze position was measured online to ensure that participants were fixating. Trials during which gaze position diverted >1.5° from fixation on the horizontal axis were excluded from further analyses.

#### Cued visual discrimination task

The first visual discrimination task was an adaptation of the Posner cueing task (Posner, 1980). Target stimuli were presented unilaterally for 200ms, on either the left or the right side of the monitor (50% left, 50% right). Prior to target presentation (-1300ms), a spatial cue that was predictive of the target stimulus location was presented for 300ms (80% cue validity). Participants were instructed to use this spatial cue to covertly shift their attention towards the cued location. Participants could respond until 1400ms after the onset of stimulus presentation by pressing one of two buttons on the keyboard (*S* for CCW tilted Gabor patches, *K* for CW). A variable inter-trial interval (ITI) randomly drawn from a uniform distribution between 250 and 350ms started directly after a response or the end of the response window if no response was given. Participants performed 560 trials per session of this task, distributed over two runs of 280 trials.

#### Uncued visual discrimination task

The other discrimination task was a classical visual discrimination task. Target stimuli were presented centrally for 200ms and there were no further manipulations. Besides indicating the orientation of the target stimulus, participants were also asked to report the confidence in their decision. Participants were instructed beforehand to distribute their confidence reports evenly, to prevent participants from only reporting low confidence answers under this challenging task. Orientation and confidence reports were simultaneously given by pressing one of four buttons on the keyboard (*A* for high confident CCW tilted Gabor patches, *S* for low confident CCW, *K* for low confident CW, *L* for high confident CW). Again, participants had a 1400ms time window to respond (response window aborted at response), followed by a 250ms-350ms ITI. In total, participants performed 600 trials of the uncued visual discrimination task per session, distributed over two runs of 300 trials.

#### Liberal and conservative visual detection task

During the liberal visual detection task, target stimuli were presented centrally for 200ms as well. The noise stimulus (a circle containing dynamic noise) was presented on every trial, but target stimuli were only shown on 50% of trials. Participants were instructed to report whether they saw a target stimulus or not, by pressing *S* (for target absent) or *K* (for target present). Target stimuli were orientated CCW and CW as in the discrimination tasks, but this orientation was not task-relevant and could thus be ignored. The response window was again 1400ms from stimulus onset, followed by a 250ms-350ms ITI. Response bias was manipulated towards more liberal answers by means of negative auditory feedback in the form of a buzzer sound after missed target stimuli (i.e., misses in signal detection theory), presented immediately after the response. Participants performed 480 trials (two runs of 240 trials) of the liberal visual detection task on each session.

The conservative detection task was like the liberal detection task, with the only difference being that response bias was manipulated towards more conservative responses. In this case, negative feedback was provided after falsely reporting the presence of a target stimulus (i.e., false alarms in signal detection theory). Again, participants performed 480 trials (two runs of 240 trials) per session of this task.

Because the buzzer sounds might have increased participants’ arousal and pupil size on the following trial, we also performed the analyses described below after excluding trials that followed a buzzer sound. With this, we removed 12.4% of trials of the liberal and conservative detection tasks (or 3.3% of all trials). Removing these trials from further analyses did not change the results (**Supplementary Figure 5**).

#### Staircasing procedure

We titrated performance on all tasks to 75% correct. To this end, the volume of the target tones or the opacity of the Gabor patches (i.e. the signal strength) was varied according to the weighted transformed up/down method proposed by Kaernbach (Kaernbach, 1991), whilst the (auditory or visual) noise was kept constant. In short, the signal strength was increased with one step after erroneous responses, and decreased with three steps after a correct response (i.e. “1-up-3-down staircase”).

Each experimental session started with a staircase procedure to find a good estimate of each subject’s auditory discrimination (session 1 and 2) or detection (session 3) threshold. Participants performed three runs of 60 trials each. We interlaced two Kaernbach staircases (30 trials each) to improve the threshold estimate. The first run started with relatively high signal strength so that participants could get acquainted with the stimuli. The second and third run started with the mean signal strength of the reversals (excluding the first reversal) of the first and second run, respectively. After finishing all staircase runs, the signal strength for the main auditory tasks that would supposedly lead to 75% performance was calculated by averaging all reversals (excluding the first reversals) across the two interlaced staircases. The auditory staircase task was similar to the main auditory tasks, but with a few differences. The staircase task had shorter inter trial intervals (1000ms) and the task required a response on each trial. Participants did not have to indicate the confidence in their decisions during the staircase task, and they were stimulated to withdraw from responding until the offset of the target sounds by flashing the fixation dot in red if they had answered too quickly.

The visual tasks were titrated to 75% performance in a similar fashion in a separate intake session (Nuiten et al., 2023).

### Eye-tracking acquisition and preprocessing

Gaze position and pupil size were measured with an EyeLink 1000 eye tracker (SR Research, Canada) during the experiment at 500Hz. Nine-point calibration was performed at the start of each run to ensure high data quality. Furthermore, we used a head-mount with chinrest to minimize participants’ head movements. Participants were instructed to move their heads as little as possible and to try to refrain from blinking during the trials.

The pupillometry data of all tasks was preprocessed in the exact same way. Pupil traces were lowpass filtered at 10Hz, blinks were linearly interpolated and the effects of blinks and saccades on pupil diameter were removed via finite impulse-response deconvolution (Knapen et al., 2016). We removed all trials for which the eyes were closed during the baseline period from further analyses.

### Data analysis

#### Analysis for data collapsed across all tasks

To assess the overall shape of the relationship between prestimulus pupil-linked arousal and perceptual decision-making, we first combined the data of all tasks (excluding all pharmacologically manipulated data). To quantify prestimulus pupil-linked arousal, we took the minimally preprocessed pupil traces of all tasks, and we calculated the average pupil size in the baseline window (-500ms – 0ms) leading up to each stimulus presentation (Figure 1B). Throughout this manuscript, we use the term pupil-linked arousal to refer to spontaneous fluctuations in arousal state as measured by pupil size (Joshi et al., 2016; Reimer et al., 2014, 2016). Note that for the cued visual discrimination task, we also took the 500ms before target presentation (not cue presentation) as the baseline window. We excluded all trials for which the eyes were closed during the baseline window, as well as trials for which pupil size during the baseline window was smaller or larger than three standard deviations from a subjects’ mean baseline pupil size. Next, for each run of each task separately, we assigned each trial to one of twenty equally sized bins based on the average prestimulus pupil size (note that we use five bins for the Analyses performed separately for each sensory modality and decision type). The binning procedure was performed per run because it is not possible to assess whether pupil size differences between runs are the result of arousal fluctuations or differences between the exact head location (even though we used a head-mount). Therefore, we are looking at arousal fluctuations that occur within experimental runs (**Figure 1C**, shown for five bins).

After having assigned all trials to twenty bins, we calculated Signal Detection Theoretic sensitivity (SDTs d’ (Green & Swets, 1966)) and average reaction time (RT) as our dependent measures of perceptual decision-making for each run. We also calculated the mean pupil size for each bin to replace (equally spaced) bin numbers with (not necessarily equally spaced) mean prestimulus pupil size values, to do justice to the true relationship between pupil size and decision behavior. Next, we subsequently averaged over runs, sessions (in the case of auditory discrimination, for which we have two sessions), and tasks, leaving us with mean d’ and RT for 20 bins for each subject. To assess the shape of the relationship between arousal and perceptual decision-making, we next used linear mixed models (see below Mixed linear models).

#### Mixed linear models

We used a mixed linear modeling approach implemented in the Python-package *Statsmodels* (Seabold & Perktold, 2010) to quantify the dependence of behavioral sensitivity and reaction time on pupil size bin (de Gee et al., 2020). Specifically, we fitted two mixed models to test whether pupil response bin predominantly exhibited a monotonic effect (first-order), or a non-monotonic effect (second-order) on the behavioral metric of interest (*y*). The fixed effects were specified as:

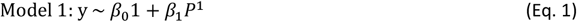

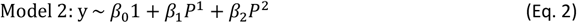

with *β* as regression coefficients and *P* as the average baseline pupil size in each bin. We included the maximal random effects structure justified by the design (Barr et al., 2013): intercepts and pupil size coefficients could vary with participant. The mixed models were fitted through restricted maximum likelihood estimation. The two models were then formally compared based on Akaike Information Criterion (AICS) (Akaike, 1974) and Bayesian Information Criterion (BIC) (Schwarz, 1978).

#### Analyses performed separately for each sensory modality and decision type

After having assessed that the shape of the overall relationship between pupil-linked arousal and perceptual decision-making was quadratically shaped, we investigated whether this relationship also held for the different sensory modalities and decision types in our dataset. We collapsed the data of all tasks over the relevant features (i.e., auditory (2 tasks), visual (4 tasks), detection (3 tasks), discrimination (3 tasks)). We next treated the data as before (see Analysis for data collapsed across all tasks), but this time we assigned the trials to five equally populated bins (instead of twenty) to compensate for the lowered power after splitting the data (Figure 1B and C). Next, we performed polynomial regression (see below Polynomial regression) to assess the shape of the relationship between arousal and perceptual decision-making in the different sensory modalities and decision types in our dataset.

#### Polynomial regression

To assess the shape of the relationship between arousal and perceptual decision-making (quadratic or linear), we performed second order polynomial regression. Because we were merely interested in the shape of the relationship, and to minimize the influence of pupil size differences between subjects on our results, we first normalized our data in the pupil dimension (i.e., essentially centering the data around a common mean). Next, we modeled the relationship between our observed behavior (d’ and RT) and prestimulus pupil size as a negative quadratic relationship, and as a (unsigned) linear relationship using ordinary least squares linear regression with freely varying intercepts. We extracted the relevant beta coefficient from each model (ß_1_ for the linear model and ß_2_ for the quadratic model) for each subject and tested whether the coefficients were significantly different from zero using one-sample t-tests (α=.05). We first tested the significance of the linear model (two-sided), followed by the quadratic model. After having established that the overall relationship between prestimulus pupil and sensitivity/RT was negatively/positively quadratically shaped, respectively, we performed one-sided tests for the quadratic beta coefficients. To further investigate the effect of pupil-linked arousal on sensitivity, we repeated the polynomial regression analyses with hit rate and false alarm rate as the dependent variables (**Supplementary Figure 4**).

#### Regressing out time-on-task

To test whether the (quadratic) relationship between pupil-linked arousal and performance could be (partly) driven by time-on-task (van den Brink et al., 2016), we linearly regressed time on task (trial number, per run) out of the pupil size data. After that, we repeated the Linear Mixed Model analysis with 20 bins and with data averaged across all tasks to test the shape of the relationship between prestimulus pupil size and perceptual decision-making after regressing out time-on-task (**Supplementary Figure 2**).

#### Regressing out evoked pupil response on the previous trial

To test whether the (quadratic) relationship between pupil-linked arousal and performance could be (partly) driven by the evoked pupil response on the previous trial, we linearly regressed the preceding evoked pupil response out of the pupil size data. The evoked pupil response was calculated as percent signal change relative to the average pupil size of a given run. The stimulus evoked pupil response was defined as the maximum pupil size in the 1 second (for the visual tasks) or 2 seconds (for the auditory tasks) after stimulus presentation as compared to the average baseline pupil size before the target (baseline: -500-0ms before target onset). After that, we repeated the Linear Mixed Model analysis with 20 bins and with data averaged across all tasks to test the shape of the relationship between pre-stimulus pupil size and perceptual decision-making after regressing out the evoked pupil response to the preceding trial (**Supplementary Figure 3**).

#### Binning analysis based on the evoked pupil response on the previous trial

To further test whether the (quadratic) relationship between pupil-linked arousal and performance could be (partly) driven by the evoked pupil response on the previous trial, we repeated the binning procedure described above (20 bins), but this time the bins were made based on the evoked pupil response on the previous trial. The stimulus evoked pupil response was again defined as described above. After this new binning procedure, Linear Mixed Model analysis was performed as before to assess the shape of the relationship between the evoked pupil response on the preceding trial and performance on the current trial (**Supplementary Figure 3**).

### Computational model

We considered a continuous-firing-rate-based, population-based model of neural dynamics to describe a generic decision-making task (Wong & Wang, 2006), which we adapt here for our detection/discrimination task. The model describes in detail the temporal evolution of the global synaptic conductances corresponding to the NMDA and GABA receptors of two competing excitatory populations and one inhibitory (PV) population (Figure 3A) The model is described by the following equations:

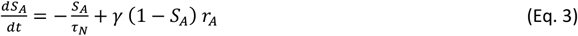

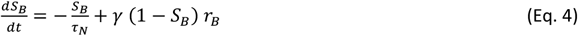

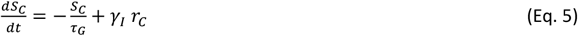

Above, S_A_ and S_B_ correspond, respectively, to the NMDA conductances of selective excitatory populations A and B, and S_C_ corresponds to the GABAergic conductance of the inhibitory population. The parameters in these equations take the following values: τ_N_=60 ms, τ_G_=5 ms, γ=1.282 and γ_I_=2. The variables r_A_, r_B_ and r_C_ are the mean firing rates of the two excitatory populations and one inhibitory population, respectively. We obtain their values by solving, at each time step, the transcendental equation *r*_*i*_ = *ϕ*_*i*_(*I*_*i*_), with *ϕ* being a transfer function of the population (specified below) and I_i_ being the total input current to population ‘i’, given by

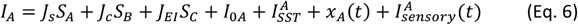

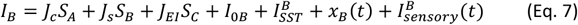

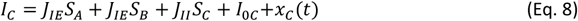

The parameters J_s_, J_c_ are the self- and cross-coupling synaptic terms between excitatory populations, respectively. J_EI_ is the coupling from the inhibitory populations to any of the excitatory ones, J_IE_ is the coupling from any of the excitatory populations to the inhibitory one, and J_II_ is the self-coupling strength of the inhibitory population. The parameters I_0i_ with i=A, B, C are background inputs to each population. Parameters in these equations take the following values: J_s_=0.49 nA, J_c_=0.0107 nA, J_IE_=0.3597 nA, J_EI_=-0.31 nA, J_II_=-0.12 nA, I_0A_=I_0B_=0.3294 nA and I_0C_=0.26 nA. Note that these last two parameters correspond to the background input that pyramidal (A,B) and PV cells (C) received from other circuits in the brain. Parameter values have been taken or slightly adapted from previous work (Mejías & Wang, 2022; Wong & Wang, 2006). The term I^i^_SST_ denotes the input to each excitatory population from its corresponding SST population (see details about this term below).

The term x_i_(t) with i=A, B, C is an Ornstein-Uhlenbeck process, which introduces some level of stochasticity in the system. It is given by

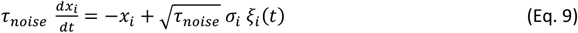

Here, ξ_i_(t) is a Gaussian white noise, the time constant is τ_noise_=2 ms and the noise strength is σ_A,B_=0.03 nA for excitatory populations and σ_C_=0 for the inhibitory one.

The last term in Eqs. 6 and 7 represents the external sensory input arriving to both populations. Assuming a detection task in which the subject has to detect the sensory stimulus A, the input is given by

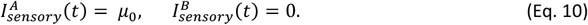

The average stimulus strength is μ_0_=0.01326 unless specified otherwise (such as in Figure 3C-D, where the performance and reaction time is studied for different stimulus strength values). The stimulus is present during the whole duration of the trial (for simulations in which multiple trials are simulated one after the other, as in Fig. 3G, the values for these currents can be switched depending on which input, A or B, is present in each trial). The discrimination task can be simply stimulated by alternating the population which receives the input in the equation above, and therefore provides equivalent results.

The transfer function ϕ_i_(t) which transform the input into firing rates takes the following form for the excitatory populations:

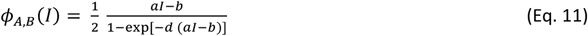

The values for the parameters are *a*=135 Hz/nA, *b*=54 Hz and *d*=0.308 s. For the inhibitory population a similar function can be used, but for convenience we choose a threshold-linear function:

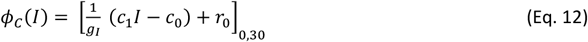

The notation [*x*]_0,30_ denotes a minimum value of zero (rectification) and a maximum value at 30 spikes/s (saturation). The values for the parameters are g_I_=4, c_1_=615 Hz/nA, c_0_=177 Hz and r_0_=5.5 spikes/s. It is sometimes useful for simulations (although not a requirement) to replace the transcendental equation *r*_*i*_ = *ϕ*_*i*_(*I*_*i*_) by its analogous differential equation, of the form

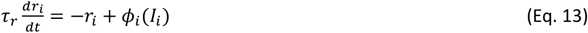

The time constant can take a typical value of τ_r_=2 ms. Note that, even though our model is only able to model continuous firing rates, the used of spike-based units such as Hertz or spikes/s is justified as our model is an extension of previous work (Wong & Wang, 2006) which is a mean-field equivalent to a spiking neural network model, and therefore it provides accurate estimations of activity mapped to spikes/s.

In addition to the two selective excitatory populations and the PV population, our model also includes the effects of top-down input mediated by selective VIP and SST populations. To introduce this, we assume that the firing rate activity of VIP and SST cells is determined, respectively, by the following equations:

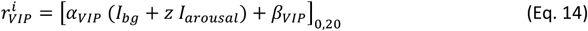

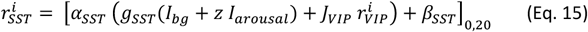

The subindex ‘i’ indicates the associated selective excitatory populations (see Figure 3A). Parameter values are α_VIP_=50, β_VIP_=0, β_SST_=20, α_SST_=32, g_SST_=2, J_VIP_=-0.1, z=0.1 and I_bg_=0.36. Note that β_SST_ and β_VIP_ correspond to the background currents received by SST and VIP cells from other circuits of the brain, and that the background current for SST is particularly high so that the elevated spontaneous activity of SST can be reproduced (C. K. Pfeffer et al., 2013). I_arousal_ indicates the arousal level of the model (assumed 0.4 in **Figure 3B-D** and variable in **Figure 3E-G**). Finally, we link the firing rate of the SST populations in Eq. 15 with the input to excitatory populations given in Eqs. 6 and 7 by assuming that:

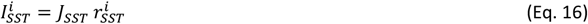

The synaptic strength is given by J_SST_=-0.001.

For each simulated trial (such as the one displayed in Figure 3B), we consider that the circuit makes a detection when the firing rate of either the excitatory population A (or B, in the case of a discrimination task) reaches a threshold of 15 spikes/s. The duration of the trial is set to T_trial_ =1.5 sec.

### Data, materials, and software availability

The data of all experiments, analysis scripts, as well as the computational modeling scripts can be found at https://osf.io/fcext/

## Notes

**Funding** This research was supported by an ERC Starting Grant from the H2020 European Research Council (ERC STG 715605 to S.v.G) and a Research Talent Grant from the Dutch Research Council (NWO; 406.17.531 to L.B. and S.v.G.).

### Competing Interest Statement

The authors have declared no competing interest.

### Summary of Updates

Main text updated, All figures updated, Supplemental figures updated.

