## Supplementary Figures for "A disinhibitory circuit mechanism explains a general principle of peak performance during mid-level arousal"

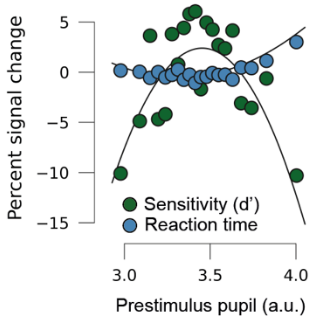


**Supplementary Figure 1. Comparison between sensitivity and reaction time effects.** Perceptual sensitivity (d’, green) and RT (blue) for small to large baseline pupils (20 bins) for all tasks combined, expressed in percent signal change with regards to the average d’ or RT across pupil bins**.** Note that the data used here is identical to the first column of Figure 2.

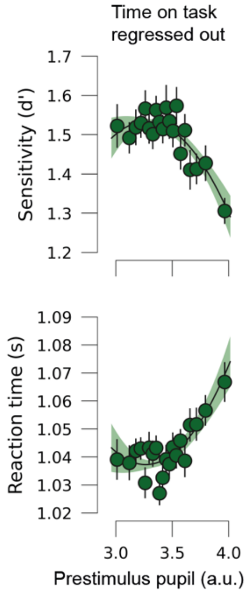


**Supplementary Figure 2. Perceptual decision-making is optimal at moderate levels of pupil-linked arousal after regressing out time-on-task.** Top row: Perceptual sensitivity (SDT's d') for small to large baseline pupils (20 bins) for all tasks combined after regressing time-on-task out of pre-stimulus pupil size. We used linear mixed effects models and formal model comparison to assess whether the relationship between pre-stimulus pupil size and sensitivity was linearly or quadratically shaped. To compare models, we calculated the difference in AIC (Akaike information criterion; ΔAIC) and BIC (Bayesian information criterion; ΔBIC) by subtracting the values for the linear model from those of the quadratic model. ΔAIC and ΔBIC were positive for sensitivity (ΔAIC=14.3, ΔBIC=10.0), suggesting a better model fit for the quadratic model. Bottom row: Same as top row but with reaction time (RT) as the dependent measure (ΔAIC=13.5, ΔBIC=9.2), again suggesting a better model fit for the quadratic model, even after regressing out time-on-task Black lines denote regression fits for the supported models.

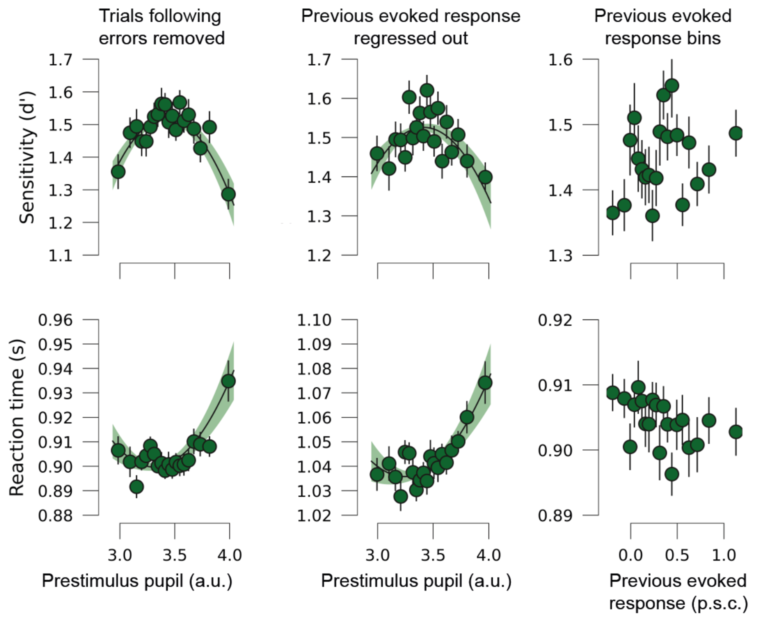


**Supplementary Figure 3. Inverted U-shaped arousal-performance relationship does not depend on significant events from the previous trial.** First column: perceptual sensitivity (SDT's d'; top row) and reaction time (RT; bottom row) for small to large baseline pupils (20 bins) after removing trials that followed errors from the analyses. We used linear mixed effects models and formal model comparison to assess whether the relationship between pre-stimulus pupil size and sensitivity was linearly or quadratically shaped after regressing out evoked responses on the previous trial. To compare models, we calculated the difference in AIC (Akaike information criterion; ∆AIC) and BIC (Bayesian information criterion; ∆BIC) by subtracting the values for the linear model from those of the quadratic model. ∆AIC and ∆BIC were positive for sensitivity and RT (d': ∆AIC=30.6, ∆BIC=26.2; RT: ∆AIC=20.5, ∆BIC=16.2), suggesting a better model fit for the quadratic model for both dependent measures of performance after removing trials following erroneous answers from the analyses. Second column: d' (top row) and RT (bottom row) for small to large baseline pupils (20 bins) for all tasks combined after regressing out the task-evoked pupil response on the previous trial of pre-stimulus pupil size on the current trial. Task-evoked pupil responses were defined as the baseline corrected maximum pupil size in the 1 (visual tasks) or 2 seconds (auditory tasks) after stimulus onset in percent signal change (p.s.c.). ∆AIC and ∆BIC were again positive for sensitivity and RT (d': ∆AIC=15.9, ∆BIC=11.6; RT: ∆AIC=13.6, ∆BIC=9.2), suggesting a better model fit for the quadratic model. Third column: d' (top row) and RT (bottom row) for small to large task-evoked pupil responses on the previous trial (20 bins). Linear mixed effects models and formal model comparison preferred neither the linear nor quadratic model for d' (∆AIC=0.0, ∆BIC=4.4), nor for RT (∆AIC=1.9, ∆BIC=6.2). Polynomial regression additionally showed that neither the linear nor the quadratic model fits were significant (all p's>.11).

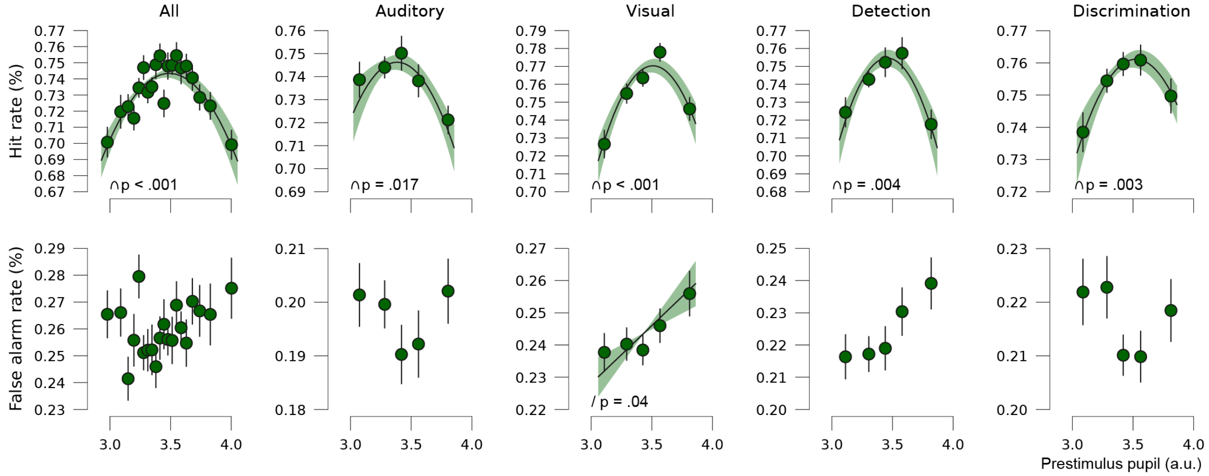


**Supplementary Figure 4. Pupil linked arousal modulates hit rate but not false alarm rate.** Top row: Hit rate for small to large baseline pupils for all tasks combined, auditory tasks, visual tasks, detection tasks, and discrimination tasks, from left to right respectively. The data was divided in twenty equally populated bins for all tasks combined and in five bins for all task variations separately. The shape of the relationship between hit rate and prestimulus pupil was assessed using polynomial regression. Solid black lines represent the average regression line of all participants with the standard error of the mean as shading. Solid black lines are shown for first and second order regression coefficients that were significantly different from zero (p<.05). Error bars represent the standard error of the mean of the within-subject variation over tasks and across pupil bins. Bottom row: as the top row, but with false alarm rate as the dependent variable.

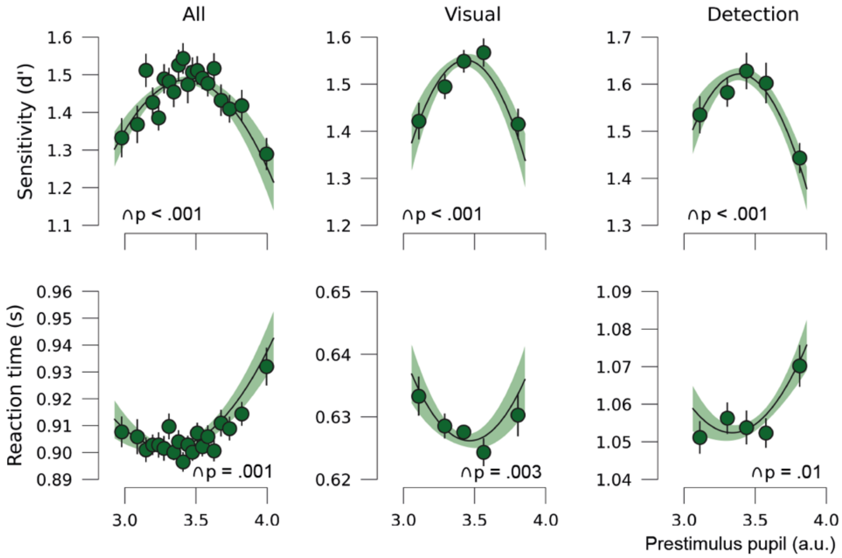


**Supplementary Figure 5. Perceptual decision-making is optimal at moderate levels of pupil-linked arousal after leaving out trials in the visual tasks that followed buzzer sounds.** Replication of Figure 2 after removing trials that followed buzzer sounds in the visual detection tasks (3.3% of all trials). Perceptual sensitivity (d', top row) and RT (bottom row) for small to large baseline pupils for all tasks combined, auditory tasks, visual tasks, detection tasks, and discrimination tasks, from left to right respectively. We show polynomial regression lines for first and second order fits when significantly different from zero (average regression line of all participants, shading reflects standard error of the mean (SEM), see Methods). Error bars on the data points represent SEM of the within-subject variation across tasks.

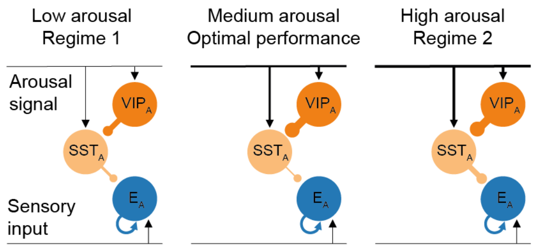


**Supplementary Figure 6. Illustration of the dynamics between VIP and SST under the influence of arousal.** When arousal is low (Regime 1), VIP activity inhibits SST firing, which disinhibits excitatory population E, resulting in increasing performance. At medium arousal levels, VIP maximally inhibits SST firing, which maximally disinhibits E, resulting in optimal performance. When arousal is high (Regime 2), VIP activity saturates, making it impossible for VIP inhibition to compensate for the excitatory effects of the arousal signal to SST. SST now starts to inhibit E, resulting in declining performance. For clarity, only a part of the model is shown here (see Figure 3A for the complete model). Thickness of the lines indicates the strength of the projections.
